# Spike mRNA Vaccine Encapsulated in a Lipid Nanoparticle Composed of Phospholipid 1,2-Dioleoyl-sn-Glycero-3-Phosphoethanolamine Induced Potent B- and T-cell Responses Associated with Protection against COVID-19 in Hamsters

**DOI:** 10.1101/2024.10.05.616797

**Authors:** Afshana Quadiri, Swayam Prakash, Latifa Zayou, Nisha Rajeswari Dhanushkodi, Amruth Chilukuri, Gemma Ryan, Kelly Wang, Hawa Vahed, Lbachir BenMohamed

## Abstract

Lipid nanoparticles (LNPs) have recently emerged as one of the most advanced vehicle platforms for efficient in vivo delivery of nucleoside-modified mRNA vaccine, particularly for COVID-19. LNPs comprise four different lipids: ionizable lipids, helper or neutral lipids, cholesterol, and lipids attached to polyethylene glycol (PEG). Studies on using the mRNA-LNP platform for vaccines have largely focused on the nucleic acid cargo with less attention to the LNP vehicle. While the LNPs protect mRNA from degradation and efficiently deliver the mRNA to antigen-presenting cells the effect of lipid composition and biophysical properties on the immunogenic and protective mRNA vaccine remain to be fully elucidated. In the present study, we used SARS-CoV-2 Spike-mRNA as a prototype vaccine, to study the effect of 4 different of LNPs with various lipid compositions. We demonstrate that when the same Spike-mRNA was delivered in the LNP4 formulation based on phospholipid 1,2-dioleoyl-sn-glycero-3- Phosphoethanolamine it outperformed the immunogenicity and protective efficacy of three LNPs (LNP1, LNP2, and LNP3) that are based on different lipids. Compared to other three LNPs, the LNP4: (*i*) enhanced phenotypic and functional maturation of dendritic cells; (*ii*) induced strong T-cell responses, (*iii*) increased secretion of proinflammatory, pro-follicular T helper (Tfh) cell cytokines; (*iv*) induced higher neutralization IgG titers; and (*v*) and provided better protection against SARS-CoV-2 infection and COVID-19 in the hamster model. We discussed the potential mechanisms by which LNP which include the phospholipid 1,2-dioleoyl-sn-glycero-3-phosphoethanolamine may activate protective B- and T-cell responses.

## INTRODUCTION

Pathogens exhibit multiple ways to evade host’s immune system. They adapt to their host during several developmental stages and express different proteins at each life stage. The later makes it complicated to identify the targets of a vaccine [1–3]. The development of safe and efficacious vaccination against diseases that cause substantial morbidity and mortality is the greatest contribution of any health intervention [4–6]. Edward Jenner observed that inoculation of pus from cowpox lesions into hosts rendered them immune to smallpox [7]. Louis Pasteur developed live-attenuated cholera, and live-attenuated rabies vaccines beginning of the modern era of immunization. Advances in understanding of viral pathogens have led to the live oral polio, measles, rubella, mumps and varicella virus vaccines[8]. The beginning of the 21^st^ century marked a revolution in molecular biology and provided insights into microbiology and immunology allowing a greater understanding of pathogen immunogens, host responses to vaccination and improved vaccine delivery [8, 9].

SARS-CoV-2 has prompted a wave of new vaccines and methods in which people gain protection from the severely challenging complications of SARS-CoV-2 [10]. Several different SARS-CoV-2 vaccine formulations were approved including mRNA and vector vaccines [11]. The main goal of vaccines is to induce high levels of neutralizing antibodies that offers protection against disease [11–13]. The use of LNPs in COVID-19 mRNA vaccines gained significant attention, for delivering the mRNA to cells as it has several advantages over viral vectors for gene therapy applications, including low cytotoxicity and immunogenicity, simple production, and great scalability [14, 15]. LNPs are spherical drug delivery vehicles with at least one lipid bilayer and an aqueous interior compartment, which provide several benefits, including formulation simplicity, self-assembly, biocompatibility, high bioavailability, and the potential to transport large payloads [16]. mRNA is inherently unstable and prone to rapid degradation by nucleases and self-hydrolysis. LNPs are efficient in mRNA encapsulation and protect it from extracellular ribonucleases and assists with intracellular mRNA delivery [17, 18]. LNPs are composed of 4 lipid components (i) ionizable lipids, which are primarily required for mRNA complexing; (ii) helper lipids, which enhance the properties of LNPs (stability and delivery efficiency); (iii) cholesterol, which provides structural stability to the LNPs (vehicle); and (iv) surface PEGylation with poly(ethylene glycol) (PEG) or PEG-derivatives, which reduces host-immune recognition and improves systemic circulation [16]. Ionizable lipids are the most essential components of LNPs, as it is neutral in the physiological pH environment but positively charged under low pH. It forms complex with the negatively charged mRNA and subsequently releases it in the cytosol, [19]. The nucleic acid encapsulated in LNPs is delivered through absorption by the cell plasma membrane and the eventual uptake into the cell by endocytosis. After endocytosis, the nucleic acid is released inside the cell. The adsorption of the LNPs is facilitated electrostatically because of the difference between the negatively charged cell membrane and the positively charged LNPs [20]. When the LNPs enter the cell, the nucleic acid is released from the cationic carrier, and the LNPs’ charge is neutralized by anionic lipids found in the cells. This neutralization of the LNPs stops electrostatic attraction between the lipids and nucleic acids, disrupting the structure of the nanoparticles and causing a non-lamellar structure thus primarily delivering mRNA through high mRNA encapsulation, greater stability, and pH-sensitive in vivo delivery [21].

Currently, there are several mRNA-based vaccines against SARS-CoV-2 which utilize different LNPs compositions, of which two are Pfizer-BioNtech’s BNT162b2 and Moderna’s mRNA-1273 [22]. Both the Pfizer-BioNtech BNT162b2 and Moderna mRNA-1273 contain mRNA encoded with the SARS-CoV-2 spike protein which when delivered into the cytoplasm, the mRNA triggers an immune response through the translation of the mRNA into the spike protein [23]. Pfizer-BioNtech uses an ionizable cationic lipid called ALC-0315 which they have licensed from Acuitas. The nanoparticles used in the Pfizer-BioNtech BNT162b2 are ALC-0315, ACL-0159, 1,2-stearoyl-sn-glycerol-3-phosphocholine (DSPC), and cholesterol. ALC-0159 is a polyethylene glycol conjugate, meaning it is a PEGylated lipid [24]. Moderna uses its own unique and proprietary ionizable cationic lipid called SM-102. The nanoparticles used in the Moderna mRNA-1273 are 1,2-stearoyl-sn-glycerol-3-phosphocholine (DSPC), polyethylene glycol 200-dimyristoyl glycerol (PEG2000-DMG), cholesterol, and SM-102 [18, 25]. In both mRNA-based vaccines, the lipids used are 1,2-stearoyl-sn-glycerol-3-phosphocholine (DSPC), cholesterol, a type of PEGylated lipid, and an ionizable cationic lipid. Modifications made to the cholesterol used in these mRNA-based vaccines increased the cellular uptake of LNPs [26]. The PEG-lipid, with its chemical structure of a hydrophilic and hydrophobic region, allows penetration across lipid membranes and influences the size of the particle [27, 28]. The ionizable lipid design has undergone numerous improvements leading to much higher transfection potency. The first ionizable lipid reported AL1 synthesized for nucleic acid delivery, exhibited low therapeutic levels of Improvements in the lipid design resulted in nearly 8000-fold improvements in the therapeutic index [29]. Introducing a ketal group in ionizable lipid led to 10% increase in gene silencing by LNPs [30].

The challenges remaining for mRNA-LNP vaccines involve the complexity associated with identifying the best formulation. The detailed mechanistic knowledge of how LNPs assist in the endocytosis of mRNA is still lacking. This makes improvements in the design and formulation of LNP slightly difficult. For most formulations, the bottleneck has not been identified, whether it is endocytosis, endosomal escape, stability of the mRNA, DC activation, or something different [31]. Much research has been done to improve endosomal escape and transfection efficiency of LNPs by modifying lipid compositions. However, compared to the extensive research done exploring different lipid formulations, the nanostructure of LNPs or the specific packing of lipids and nucleic acids into the LNP has not been fully investigated. Over the past 20 years, ionizable lipid design has undergone numerous improvements, the first ionizable lipid reported AL1 also known as DODAP, didn’t provide significant therapeutic levels of delivery unless high doses were given. Improvements to the lipid design resulted in nearly 8000-fold improvements in the therapeutic index[32]. Interestingly, the ester-containing analogue, was found to be ineffective while a ketal-containing compound exhibited 10-fold more activity than the effective dose required to achieve 50% gene silencing [26, 32]. LNP size plays a critical role in delivery for larger nanoparticles, however, research is required to understand whether and how LNP size influences mRNA delivery. A comparative evaluation of different LNPs is, therefore, required to determine the optimal parameters for the desired vaccination. In the present study, we have investigated the pre-clinical efficacy of four different LNPs. The 4 LNPs (described in this study as LNP1, LNP2, LNP3, LNP4) differed in their phospholipid compositions or ionizable lipid relative amount, which we found to significantly affect the degree of protection from COVID-19 disease.

In this study, we observed that LNP4 that contain a phospholipid constituent named as 1,2- dioleoyl-sn-glycero-3-Phosphoethanolamine (DOPE) has higher potency at inducing better immune responses and better protection in hamsters upon SARS-CoV-2 infection. Compared to other 3 LNPs, the LNP4: (*i*) enhanced phenotypic and functional maturation of dendritic cells; (*ii*) induced strong T-cell responses, (*iii*) increased secretion of proinflammatory cytokines, pro- follicular T helper (Tfh) cell cytokines; (*iv*) induced higher neutralization IgG titers and (*v*) and provided better protection against SARS-CoV-2 infection and COVID-19 in the hamster model. Potential mechanisms by which LNP that include phospholipid 1,2-dioleoyl-sn-glycero-3-phosphoethanolamine activate protective B and T cells are discussed. The findings point to the need for understanding and adjusting lipid formulations to achieve higher efficacy of LNP-mRNA-based vaccine.

## MATERIALS AND METHODS

### LNP formulation

Four Customized GenVoy-ILM based lipid nanoparticles containing a common ionizable cationic lipid and with differing compositions of phospholipid, cholesterol and stabilizer were provided by Precision NanoSystems (PNI) (currently Cytiva) for this project. The 4 LNPs differed in their phospholipid compositions or ionizable lipid relative amount. LNPs with these compositions are available for purchase from PNI upon request using the identifiers IL00V02, IL00V03, IL00V29, and IL00V41. For the convenience of this study, we have used the nomenclature as LNP1, LNP2, LNP3, and LNP4 to identify IL00V02, IL00V03, IL00V29, and IL00V41 respectively. Each of these 4 LNPs comprised of a unique phospholipid that are the second listed lipid component of the LNP. Different phospholipids are associated with the 4 LNPs as follows: (***i***) LNP1 (IL00V02) and LNP3 (IL00V29) comprised of DSPC (1,2-distearoyl-sn-glycero-3-phosphocholine), (***ii***) LNP2 (IL00V03) comprised of DOPC (1,2-dioleoyl-sn-glycero-3-phosphocholine), and (***iii***) LNP4 (IL00V41) comprised of DOPE (1,2-dioleoyl-sn-glycero-3-phosphoethanolamine). While the ionizable phospholipid that was used in all the formulations is the same, the amount used between formulations varied. LNP3 (IL00V29) and LNP4 (IL00V41) both have a “higher” amount of ionizable lipid, but the phospholipid composition is different. The physical properties associated with the four LNPs are further described in **Table 1**.

To prepare the lipid encapsulation, first the mRNA stock was diluted in PNI’s formulation buffer. Subsequently the 4 GenVoy-ILM-based lipid mixes were prepared in ethanol at 12.5 mM concentration. The LNP formulations were prepared on the NanoAssemblr Ignite using a specific formulation scheme which included a Flow rate ratio (FRR) for mRNA solution to lipid solution at 3:1; Total flow rate (TFR) at 12mL/min; Start waste at 1mL; and End waste at 0.05mL. After formulation, the post chip formulations were diluted to a total of 4X volume in sterile Ca^2+^ and Mg^2+^ free PBS. Amicon® Centrifugal Filters with a MWCO of 30 KDa (LNP1, LNP2, LNP3) or a MWCO of 100 KDa (LNP4) were used to concentrate formulations and perform ethanol removal and buffer exchange to 1X PBS (pH 7-7.3, Mg^2+^/Ca^2+^ -free). mRNA LNPs were filtered manually through a 0.22 μm syringe filter using aseptic techniques and were stored at 4°C. Finally, the particle size and polydispersity index (PDI) were analyzed using Dynamic light scattering (DLS). RNA encapsulation efficiency and concentrations were determined using a RiboGreen plate-based assay.

### Human Samples

Blood samples were obtained from healthy donors. Consenting adults were screened using a questionnaire determining their demographic information, medication usage, and comorbidities. Participants were excluded from any acquired immunodeficiency or immunomodulating medications (such as steroids or chemotherapy), pregnancy, history of cancer, history of cirrhosis or renal failure, or antibiotic use within 2 weeks of recruitment. Blood samples were taken from individuals aged 18-75. The experiments were approved by the Institutional Biosafety Committee at the University of California, Irvine (Protocol number BUA-R112). Written informed consent was obtained from all patients before inclusion.

### PBMC isolation

The blood of anonymous healthy donors was diluted with RPMI1640. Peripheral blood mononuclear cells (PBMCs) were isolated by density centrifugation using percoll. After centrifugation, the interphase containing PBMCs was collected and washed twice with RPMI1640. The cells were cultured in tissue culture dishes and incubated at 37°C, 5% CO2.

### Generation of human monocyte-derived dendritic cells (MoDCs)

The DCs in PBMC were allowed to adhere to the culture dish. On the next day, the medium was replaced with DC-medium containing 800 U/ml GM-CSF and 50 U/ml IL4. On day 3 and day 5, a total of 4 ml fresh DC-medium supplemented with 800 U/ml GM-CSF and 50 U/ml IL4 was added, respectively. On day 7, immature as well as mature DCs were harvested. As moDCs adhere loosely to the culture dish, the use of EDTA or cell scrapers is not needed. Thus, the supernatant was flushed a couple of times to obtain moDCs. Afterward, moDCs were treated with different LNPs for 24-36 hours. The cells are harvested and stained for FACS analysis. The supernatant was collected and analyzed for different cytokine

### Flow cytometry

The following anti-human antibodies were used for the flow cytometry assays: CD11c (clone BLY6 – BD BioSciences), HLA-DR (clone G46-6 – BDBioSciences), CD40 (clone 5C3 –Invitrogen), CD86 (clone 2331 (FUN-1) – BD BioSciences), and CD83 (clone HB15e – BDBioSciences). For surface stain, mAbs against cell markers were added to a total of 1 x 10^6^ cells in 1X PBS containing 1% FBS and 0.1% sodium azide (FACS buffer) for 30 minutes at 4°C. The cells were washed with FACS Buffer and finally suspended in 200ul of FACS buffer. For each sample, 50,000 total events were acquired on the BD LSRII. Ab capture beads (BD Biosciences) were used as individual compensation tubes for each fluorophore in the experiment. We used fluorescence minus controls for each fluorophore to define positive and negative populations when initially developing staining protocols. Data analysis was performed using FlowJo. Statistical analyses were done using GraphPad Prism version 7.

### Hamster mRNA-LNP immunization and SARS-CoV-2 virus challenge

The mRNA-LNP vaccines were evaluated in the outbred golden Syrian hamster model for protection against SARS-CoV- 2 USA/WA1/2020 strain. The Institutional Animal Care and Use Committee approved animal model usage experiments at the University of California, Irvine (Protocol number AUP-22-086). The recommendations in the Guide for the Care and Use of Laboratory Animals of the National Institutes of Health performed animal experiments. The sample size for each animal study was 5 (*n* = 5 per group). 6–8-week-old male Golden Syrian Hamsters (N = 45), divided into nine experimental groups were vaccinated with 1µg and 3µg of LNP1, LNP2, LNP3, and LNP4 encapsulating Spike protein intramuscularly in 100 μL of doses into the posterior thigh muscle, twice (On Day 0 and Day 21). 4 weeks after the first immunization blood sample was collected from hamsters under isoflurane anesthesia and spun at 2000g for 10 min to obtain serum. The hamsters were challenged intranasally with 100µl of 1 x 10^5^ PFU of SARS-CoV-2 USA/WA1/2020 strain diluted in sterile Dulbecco’s Modified Eagle’s media (DMEM). Weights were recorded daily until 14 days p.i. Oropharyngeal swabs were collected on day 2, 6, 10, 14 in 500µl of RNA later for virus titrations. The animals in each group were monitored daily for signs of disease and weighed until 14 days p.i.

### QuantiFERON assay

The whole heparinized blood sample was collected from each hamster for performing QuantiFERON assay to determine the secretion of IFN-γ (QuantiFERON SARS-CoV-2, Qiagen). After overnight incubation with SARS-CoV2-specific peptides, supernatant was collected from LNP-stimulated samples in vitro at 12 and 24-hour time points for IFN-γ analysis. The cytokine analysis was done on 96 well-flat bottoms. ELISA plates coated with specific capture antibodies. The supernatant obtained from LNPs stimulated moDCs was tested without any dilutions. For hamster ELISA, the sera were collected from blood isolated from immunized hamsters. ELISA was performed on sterile 96-well flat-bottom microplates coated with the Spike antigen in coating buffer overnight at 4°C. The reaction was terminated by adding 1M H2SO4. The absorbance was measured at 450 nm.

### Enzyme-linked immunosorbent assay (ELISA)

Serum antibodies for SARS-CoV-2 proteins were detected by ELISA. 96-well plates (Dynex Technologies, Chantilly, VA) were coated with 100 ng S protein per well at 4°C overnight, and then washed three times with PBS and blocked with 3% BSA (in 0.1% PBST) for 2 h at 37°C [6, 33, 34]. After blocking, the plates were incubated with serial dilutions of the sera (100 μl/well, in two-fold dilution) for 2 hours at 37°C. The bound serum antibodies were detected with HRP-conjugated goat anti-mouse IgG and chromogenic substrate TMB (ThermoFisher, Waltham, MA). The cut-off for seropositivity was set as the mean value plus three standard deviations (3SD) in HBc-S control sera.

### Neutralization Assay

Serum neutralizing activity was examined, as previously reported in [34–36]. The assays were performed with Vero E6 cells (ATCC, CRL-1586) using the SARS-CoV-2 wild-type or Delta strains. Briefly, serum samples were heat-inactivated and three-fold serially diluted (initial dilution, 1:10), followed by incubation with 100 pfu of wild-type SARS-CoV-2 (USA-WA1/2020) strain for 1 hour at 37°C. The serum-virus mixtures were placed onto Vero E6 cell monolayer in 96-well plates for incubation for 1 hour at 37°C. The plates were washed with DMEM, and the monolayer cells were overlaid with 200 μl minimum essential medium (MEM) containing 1% (w/v) of methylcellulose, 2% FBS, and 1% penicillin-streptomycin. Cells were then incubated for 24 hours at 37°C. Vero E6 monolayers were washed with PBS and fixed with 250 μl of pre-chilled 4% formaldehyde for 30 min at room temperature, followed by aspiration removal of the formaldehyde solution and twice PBS wash. The cells were permeabilized using 0.3% (wt/vol) hydrogen peroxide in water. The cells were blocked using 5% non-fat dried milk followed by the addition of 100 μl/well of diluted anti-SARS-CoV-2 antibody (1:1000) to all wells on the microplates for 1-2 hours at RT. This was followed by the addition of diluted anti-rabbit IgG conjugate (1/2,000) for 1 hour at RT. The plate was washed and developed by the addition of TrueBlue substrate, and the foci were counted using an ImmunoSpot analyzer. Each serum sample was tested in duplicates.

### Histology of animal lungs

Hamster lungs were preserved in 10% neutral buffered formalin for 48 hours before transferring to 70% ethanol. The tissue sections were then embedded in paraffin blocks and sectioned at 8 μm thickness. Slides were deparaffinized and rehydrated before staining for hematoxylin and eosin for routine immunopathology as described before [13, 34].

### Statistical analysis

Data for each assay were compared by ANOVA and Student’s *t*-test using GraphPad Prism version 7 (La Jolla, CA). As we previously described, differences between the groups were identified by ANOVA and multiple comparison procedures [37, 38]. Data are expressed as the mean ± SD. Results were considered statistically significant at a *P* value of ≤ 0.05.

## RESULTS

### 1. Lipid nanoparticles of different composition vary in their effect on the maturation and activation of dendritic cells

Much of the immunobiology of Dendritic cells (DCs) orchestrates around their role as antigen-presenting cells and in activation of T-cells. CD40 is upregulated on activated DCs and CD40L is expressed on activated T cells. Engagement of CD40 on DCs with CD40L induces positive signaling that leads to the expression of CD83/86 and the production of IL-12 in DCs.

We observed a similar phenomenon wherein the treatment of DCs with different LNPs leads to the activation and maturation of DCs. Monocytes isolated from healthy participant PBMCs (n = 4; age range 24–75 yrs) were treated with GM-CSF/IL-4 for 6 days. On day 7, immature as well as mature DCs were harvested, followed by treatment with a dose of 1.2 μg/mL of 4 different LNPs for 24 h. We assessed the frequency of surface costimulatory and HLA marker-expressing cells in LNP-treated MDDCs compared to unstimulated cells after 24 h. We found that all LNPs significantly up-regulated the percentage of IL12 and CD40-positive human MDDCs compared to unstimulated control (**Figs. 1A** and **1B**). However, the upregulation was relatively higher in cultures treated with LNP-4 (**Figs. 1A** and **1B**). We also observed a significant upregulation in the maturation markers of MDDCs -CD83 and CD86 compared to unstimulated control (**Figs. 1C** and **1D**). However, the upregulation was relatively higher in cultures treated with LNP-4 (**Figs. 1C** and **1D**).

**Figure 1.**
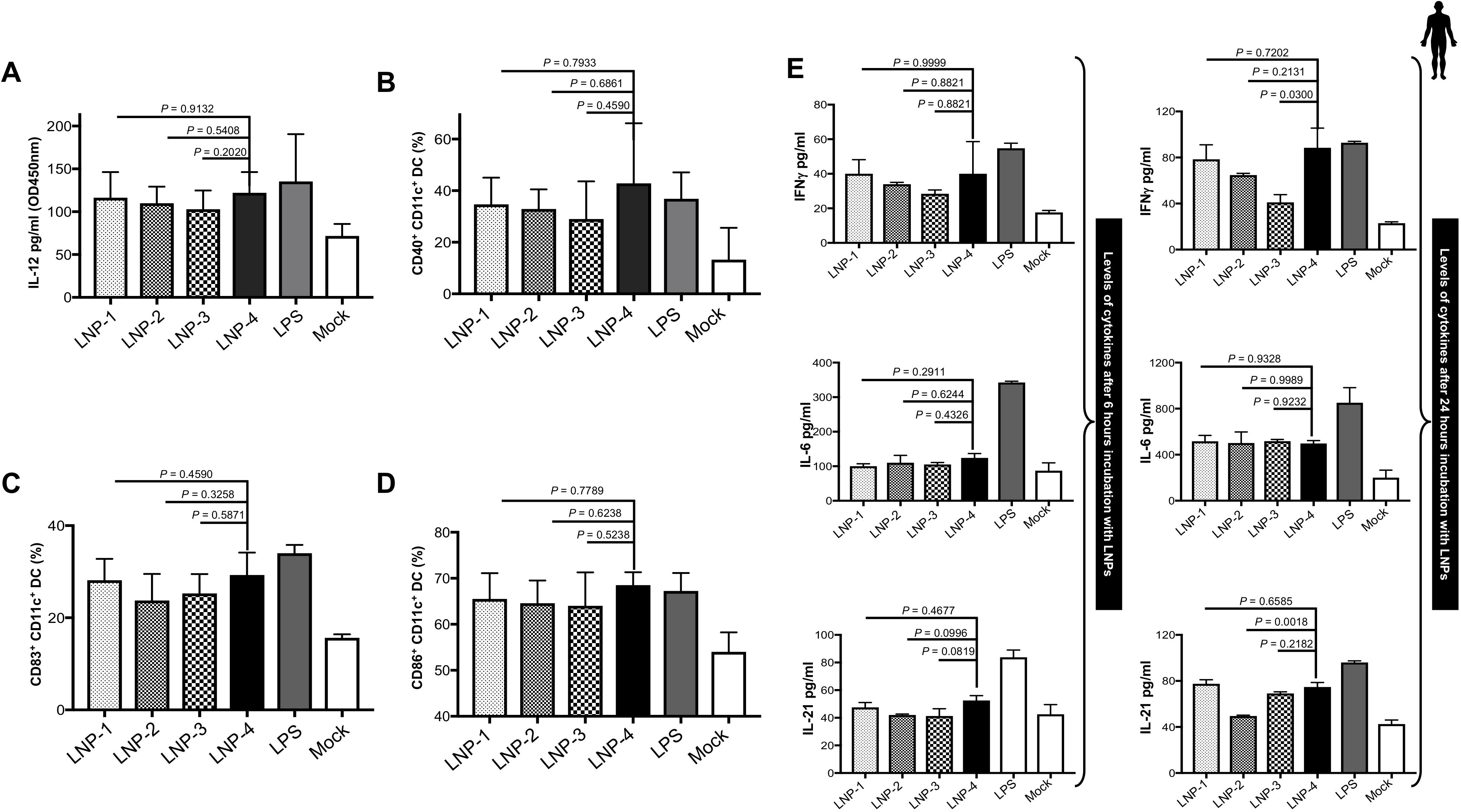
LNP stimulation and immunogenicity: **(A)** IL-12 secretion by monocyte-derived DCs after 24 h of stimulation with 1.2 μg/mL of different LNPs. (**B)** Activation status of monocyte-derived DCs as measured by CD40 expression after 24 h of stimulation with 1.2 μg/mL of different LNPs. **(C&D)** Expression of maturation markers CD83 **(C)** and CD86 **(D)** on DCs following stimulation with 1.2 μg/mL of different LNPs. **(E)** Dynamics of cytokines from PBMCS at 6, and 24 h after LNP stimulation.

Next, we tested the production of cytokines following in vitro DC maturation after 6 hours and 24 hours of incubation with different LNPs. IL-6, IL-21, and IFN-γ were markedly found to be increased after 24 h in cultures stimulated with LNPs and this increase was relatively more pronounced in cultures stimulated with LNP-4 (**Fig. 1E**). LNPs can induce pro-TFH cytokines as well as key cytokines that are efficient in activating innate immune responses like IFN-γ and LNP-4 displayed higher ability compared to other LNPs.

### 2. Vaccination with mRNA-Spike encapsulated in LNP-4 demonstrated better protection against SARS-CoV-2

We sought to assess the in vivo protection efficacy of mRNA-Spike-LNPs in the well-established SARS-CoV-2 hamster model using the Washington strain. Groups of 8-week-old Syrian golden hamsters were i.m. administrated with two doses of different (1µg and 3µg) concentrations of mRNA-Spike-LNPs, followed by intranasal (i.n.) challenge with a sub-lethal dose (1 x 10^5^ pfu) of SARS-CoV-2 Delta variant at 14-day post 2^nd^ immunization (**Fig. 2A**). Weight loss analysis showed that all hamsters in the mock group lost body weight and displayed a mean reduction of 4%- 5% of body weight by day 4 (**Figs. 2B** and **2D**). While immunized hamsters showed a mean increase of 2-3% (1µg) and 2-4% (3µg) in body weight (**Figs. 2B** and **2D**). However, hamsters immunized with LNP-4 demonstrated better weight gain and overall maximum weight (**Figs. 2C** and **2E**).

**Figure 2.**
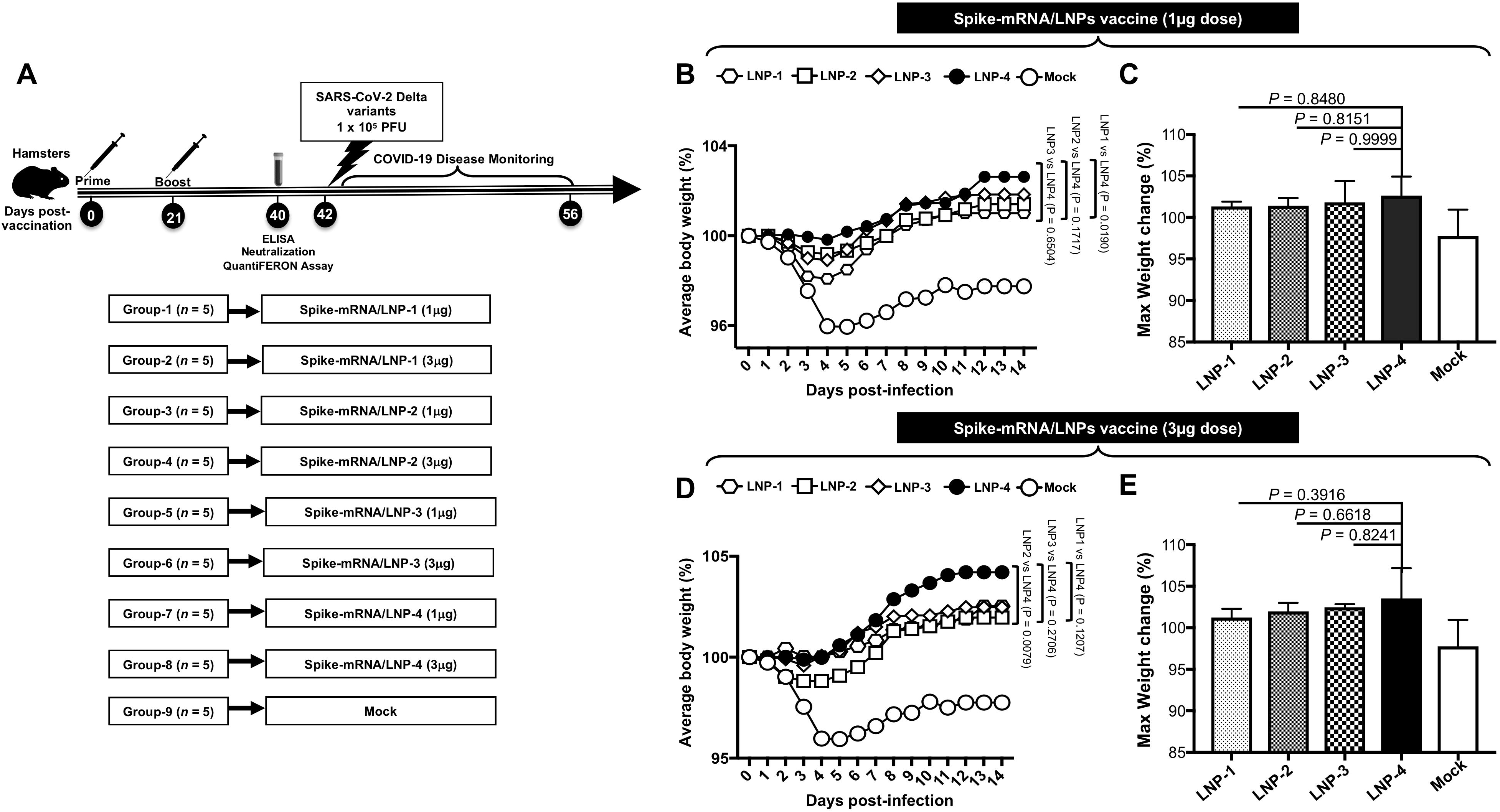
Vaccination of Syrian hamster using *mRNA-Spike encapsulated in different LNPs* followed by infection with Washington variant: **(A)** Schematic of experimental design. Syrian golden hamsters were vaccinated with 1μg and 3μg of LNP1-4 encapsulating Spike protein intramuscularly twice (21 days apart). 4 weeks after the first immunization blood sample was collected from hamsters under isoflurane anesthesia and spun to obtain serum. The hamsters were challenged with 1X10^5^ viral particles of Washington strain diluted in sterile Dulbecco’s Modified Eagle’s media (DMEM). Weights were recorded daily until 14 DPI. **(B)** Mean body weight changes following immunization (1 μg dose) and infection of hamsters with SARS-CoV-2 Delta variant, along with weight change in uninfected control hamsters. **(C)** Mean of weight gain in hamsters at 14 DPI following immunization (1 μg dose) and infection with SARS-CoV-2 Delta variant. **(D)** Mean body weight changes following immunization (3μg dose) and infection of hamsters with SARS-CoV-2 Delta variant, along with weight change in uninfected control hamsters. **(E)** Mean of weight gain in hamsters at 14 DPI following immunization (3 μg dose) and infection with SARS-CoV-2 Delta variant.

### 3. Vaccination with mRNA-Spike encapsulated in LNP-4 induces high IgG titer and neutralizing antibodies

To assess the immunogenic potential of mRNA-LNP vaccines formulated with 4 different LNP compositions, Syrian golden hamsters (n = 5 per group) were vaccinated three weeks apart with 2 doses of either mRNA-LNP vaccines i.m. Immunogenicity was assessed at approximately 1 week after dose 2 (Day 28); Spike-specific serum immunoglobulin (Ig) G or A binding antibody responses were measured by enzyme-linked immunosorbent assay (ELISA) and serum neutralizing antibody titers were measured by a plaque reduction neutralization test (PRNT). Spike mRNA encapsulated in LNP-4 immunization elicited higher antibody titers followed by LNP 3, LNP-2, and LNP- 1 (**Figs. 3A and 3D**). LNP-1 displayed lower Spike-specific binding titers. In addition to S-specific binding titers, neutralizing antibody responses in sera were evaluated. LNP-4 displayed better neutralizing potential followed by LNP-1 and LNP-2. LNP-3 displayed lower neutralizing potential (**Figs. 3B and 3E**).

**Figure 3.**
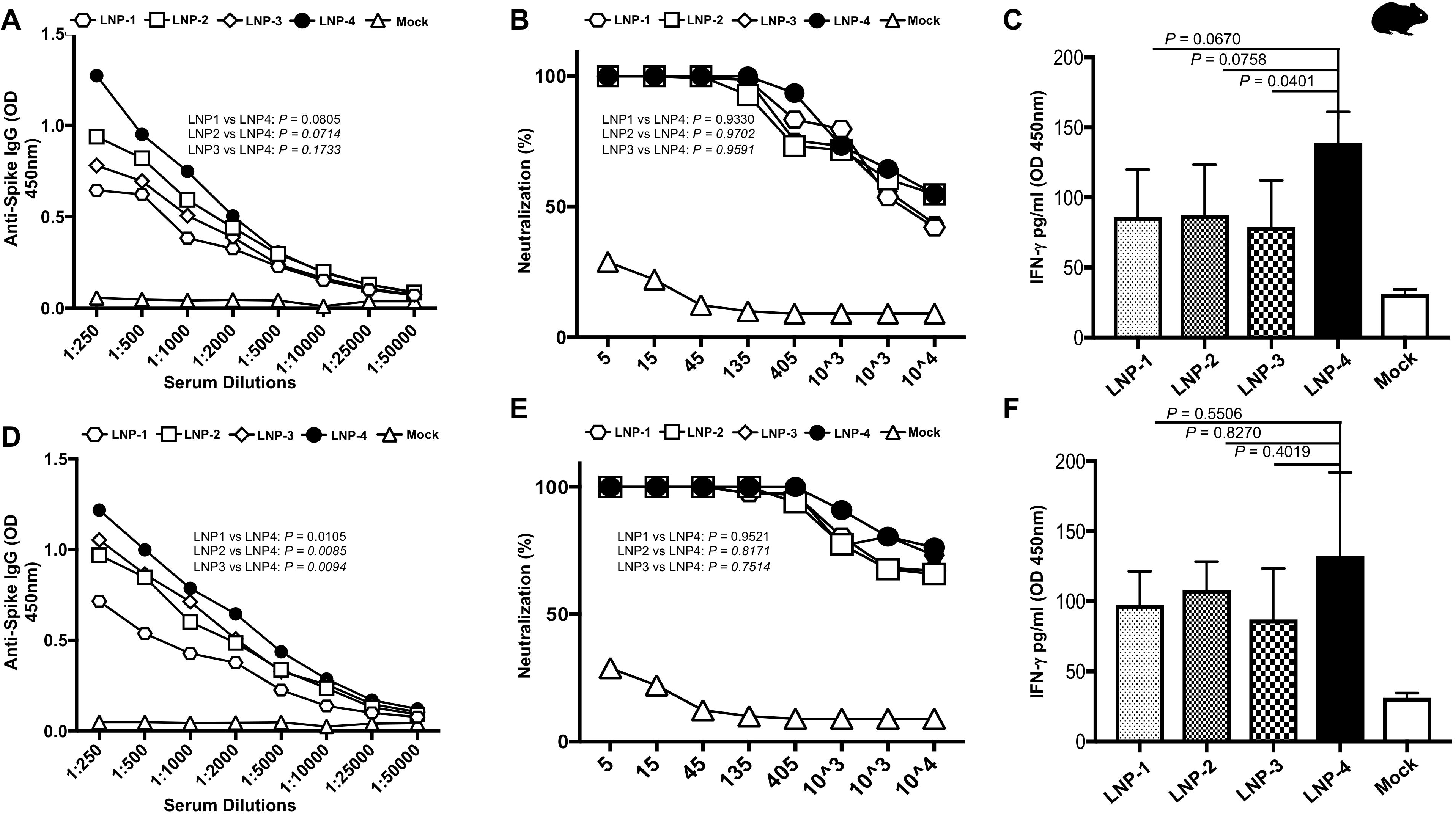
SARS- CoV2 specific immune analysis by ELISA, Neutralization and QuantiFERON assays for select LNP formulations. **(A)** Spike specific serum immunoglobulin (Ig) G or A binding antibody responses obtained from Hamsters immunized with 1 μg of Spike-mRNA/LNP as measured by enzyme-linked immunosorbent assay (ELISA). **(B)** Serum neutralizing antibody titers obtained from Hamsters immunized with 1 μg of Spike-mRNA/LNP as measured by a plaque reduction neutralization test. **C)** QuantiFERON assay to determine induction of IFN-γ cytokine in peripheral blood of hamsters immunized with 1μg of Spike-mRNA/LNP upon stimulation with Spike peptides. **(D)** Spike specific serum immunoglobulin (Ig) G or A binding antibody responses obtained from Hamsters immunized with 3μg of Spike-mRNA/LNP as measured by enzyme-linked immunosorbent assay (ELISA). **(E)** Serum neutralizing antibody titers obtained from Hamsters immunized with 1 μg of Spike- mRNA/LNP as measured by a plaque reduction neutralization test. **(F)** QuantiFERON assay to determine induction of IFN-γ cytokine in peripheral blood of hamsters immunized with 3 μg of Spike- mRNA/LNP upon stimulation with Spike peptides.

### 4. Vaccination with mRNA-Spike encapsulated in LNP-4 induced higher level of Interferon-γ

The heparinized fresh whole blood collected from each hamster immunized with different LNPs was incubated with SARS-CoV2 Spike specific peptides for 24 hours. The presence of IFN-γ in the plasma samples was determined by ELISA. Peripheral blood obtained from Hamsters immunized with LNP-4 showed higher production of IFN-γ in vitro followed by hamsters immunized with LNP-2, LNP-3, and LNP-1 (**Figs. 3C and 3F**).

### 5. Vaccination with mRNA-Spike encapsulated in LNP-4 confers protection against SARS-CoV-2 Delta variants

We next determined the protective efficacy mRNA-LNP vaccines that incorporate the Spike antigen against the highly pathogenic Delta variant (B.1.617.2). Hematoxylin and eosin staining of lung sections at day 14 p.i. showed a significant reduction in COVID-19-related lung pathology in the hamsters vaccinated with mRNA-LNP vaccines and LNP-4 compared to mock-vaccinated mice (**Fig. 4A**). This reduction in lung pathology was observed at a higher level for hamsters vaccinated with 3ug followed by hamsters vaccinated with 1ug dose of the LNP4 composition of the COVID-19 vaccine. (**Fig. 4**). Taken together, these results indicate that vaccination with our novel COVID-19 vaccine formulated with LNP-4 that constitutes Phospholipid DOPE, reduced COVID-19- related lung pathology following infection with pathogenic SARS-CoV-2 Delta variant.

**Figure 4.**
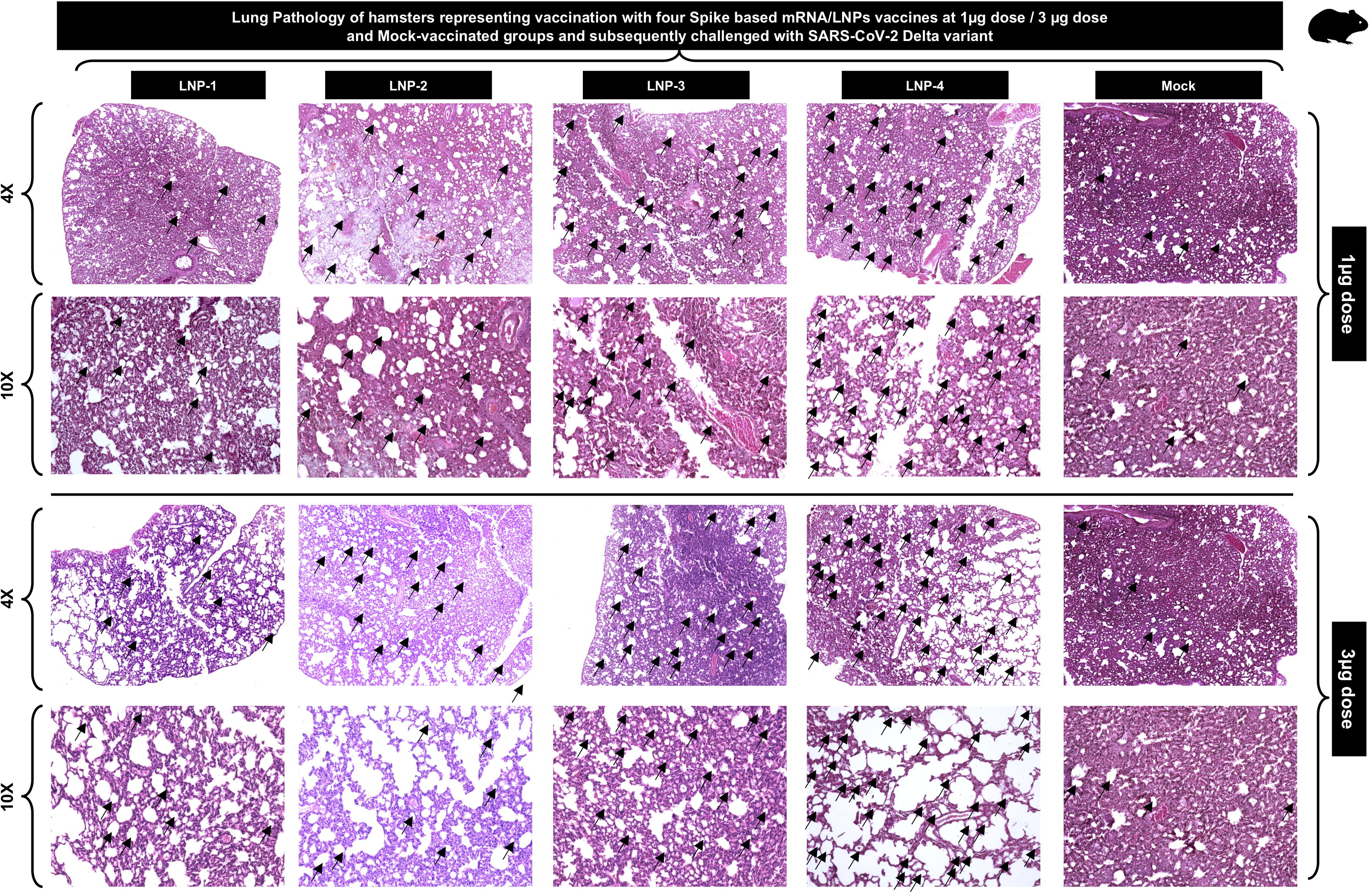
Histopathology and immunohistochemistry of the lungs from vaccinated and mock-vaccinated golden hamsters show reduced lung pathology in LNP-4 mRNA-LNP based vaccinated hamsters. Representative images of Hematoxylin and Eosin (H & E) staining of the lungs harvested on day 14 p.i. from golden hamsters vaccinated with our novel COVID-19 vaccine formulated in different LNPs (LNP-1, LNP-2, LNP-3, LNP-4) and mock-vaccinated hamsters. Hamsters were either vaccinated with individual mRNA-LNP COVID-19 vaccines at 1ug or 3ug doses. Reduced lung pathology is indicated with more number of lung vacuoles (marked with black arrows) and less degree of hemorrhage. Images were captured at 4X and 10X resolution.

### 6. Vaccination with mRNA-Spike encapsulated in LNP-4 induced strong B and T-cell responses compared to mRNA-Spike encapsulated in LNP-O

To assess the immunogenic potential of LNP-4 in comparison to LNP-O, mRNA-Spike was formulated in two different LNP compositions. Syrian golden hamsters (n = 6 per group) were vaccinated three weeks apart with 2 doses of either mRNA-LNP vaccines i.m. (**Fig. 5A**). Immunogenicity was assessed at approximately 1 week after dose 2 (Day 28); Spike-specific serum immunoglobulin (Ig) G or A binding antibody responses were measured by enzyme-linked immunosorbent assay (ELISA) and serum neutralizing antibody titers were measured by a plaque reduction neutralization test (PRNT). Spike mRNA encapsulated in LNP-4 immunization elicited higher antibody titers compared to LNP-O (**Fig. 5B**). LNP-O displayed lower S-specific binding titers. In addition to S-specific binding titers, neutralizing antibody responses in sera were evaluated. LNP-4 displayed slightly better neutralizing potential compared to LNP-O (**Fig. 5C**). LNP-O displayed slightly lower neutralizing potential. The heparinized fresh whole blood collected from each hamster immunized with different LNP-4 vs LNP-O was analyzed for T-cell responses. LNP-4 immunized blood showed better overall T-cell responses as well as display of activation markers on CD4^+^ and CD8^+^ T-cells (**Fig. 5D**).

**Figure 5.**
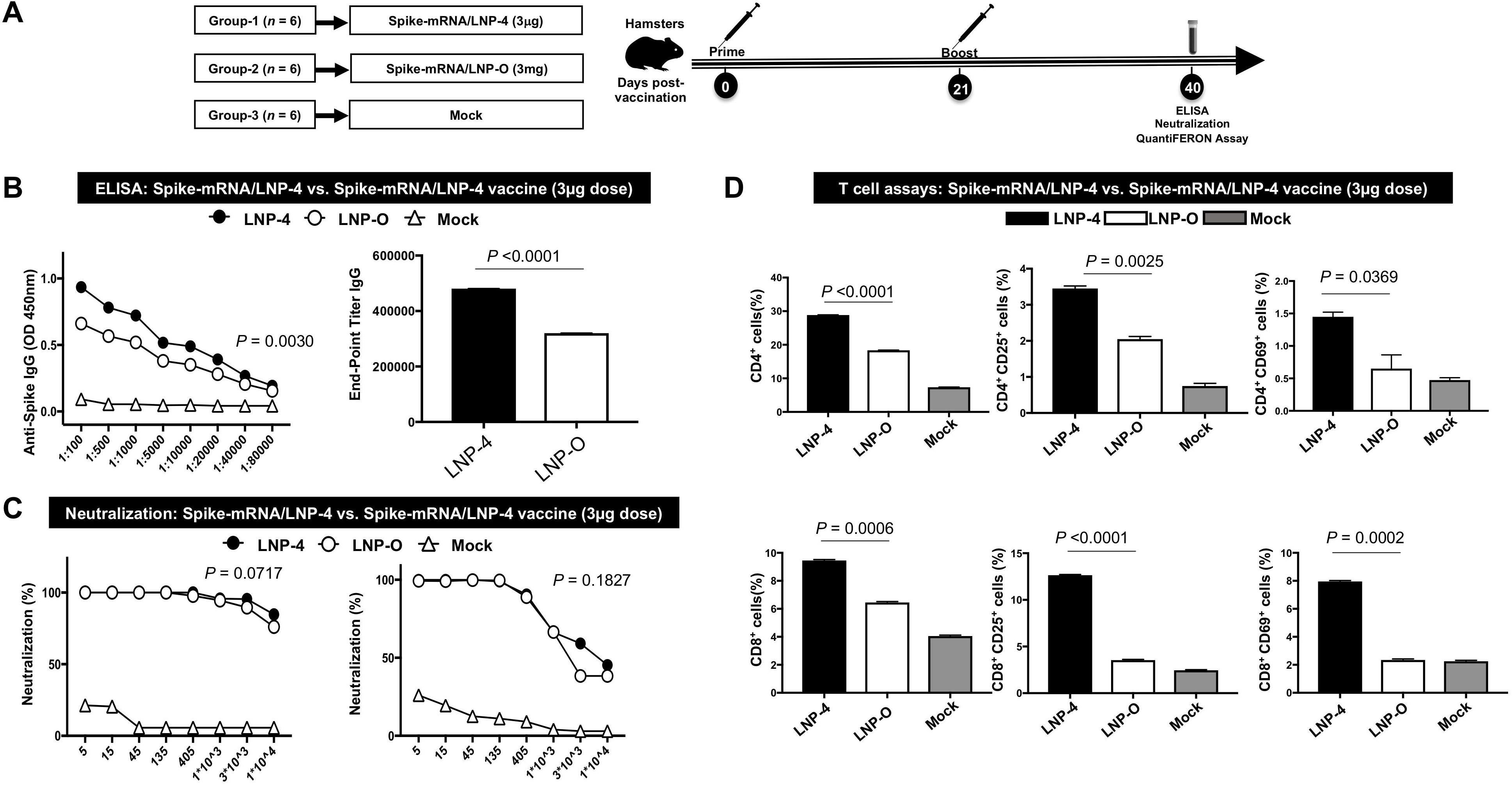
Immunogenicity comparison of LNP-4 with Off the shelf LNP (LNP-O): **(A)** Schematic of experimental design. Syrian golden hamsters were vaccinated with 3 μg of LNP4 and LNP-O encapsulating Spike protein intramuscularly twice (21 days apart). 4 weeks after the first immunization blood sample was collected from hamsters under isoflurane anesthesia and spun to obtain serum. **(B)** Spike specific serum immunoglobulin (Ig) G or A binding antibody responses obtained from Hamsters immunized with 3ug of Spike-mRNA/LNP (LNP-4 vs LNP-O) as measured by ELISA. **(C)** Serum neutralizing antibody titers obtained from Hamsters immunized with 3 μg of Spike-mRNA/LNP (LNP-4 vs LNP-O) as measured by a plaque reduction neutralization test. **(D)** Analysis of T-cell responses in the peripheral blood of Hamsters immunized with 3μg of Spike-mRNA/LNP (LNP-4 vs LNP-O).

## DISCUSSION

The need for mRNA encapsulation arises from the fragility of mRNA as a standalone. The mRNA molecule is inherently unstable and can be easily degraded by various nucleases present in the extracellular environment, and by cellular enzymes present inside the cell. When mRNA is injected into the body, the immune system perceives it as a foreign substance and tries to eliminate it. This happens very quickly, even before the mRNA has a chance to reach the target cells where it can be translated into protein. One of the main reasons why current mRNA-based vaccines targeted towards SARS-CoV- 2 need to be stored in extremely cold temperatures is to limit the degradation and chemical reactions that occur to the modified mRNA. However, encapsulating mRNA in lipid nanoparticles (LNPs) helps protect it from degradation and immune recognition, enabling it to reach the target cells and be translated into protein. The LNPs in mRNA-based vaccines are designed to fuse with the cell membrane, allowing the encapsulated mRNA to enter the cytoplasm. The ribosomes then use the mRNA as a template to synthesize the viral protein, which is subsequently presented on the cell surface to trigger an immune response. By encapsulating the mRNA in LNPs, the vaccine can achieve higher levels of protein expression and a more robust immune response than if the mRNA were injected on its own. The use of LNPs helps to prolong the duration of protein expression, allowing the immune system more time to recognize and respond to the viral protein [39].

The composition of lipid nanoparticles makes them novel compared to previous lipid-based nucleic acid delivery systems. LNPs typically consist of a lipid core matrix that solubilizes the mRNA, as well as stabilizing surfactants or emulsifiers. The lipid core matrix of LNPs is usually composed of a mixture of lipids, including phospholipids, cholesterol, and other lipids such as triglycerides, diglycerides, monoglycerides, and fatty acids [40]. These lipids are chosen for their ability to form stable particles and solubilize hydrophobic mRNA molecules. The lipid composition can be tailored to optimize delivery efficiency, stability, and safety. In the case of mRNA vaccines for SARS-CoV-2, four types of lipids are used. Namely, an ionizable cationic lipid whose positive charge can bind to the negatively charged mRNA, a PEGylated lipid meant for stability, and a phospholipid along with cholesterol designed for structural support. These components facilitate the LNPs containing mRNA to enter a cell through an endosome. When the LNP is inside the acidic endosome, the ionizable lipids become positively charged and help release the LNP and modified mRNA into the cell’s cytoplasm. The modified mRNA, now free, is translated by the ribosomes to make proteins.

Immunization of mice with self-amplifying RNA encoding the virus spike protein encapsulated in LNP formulation produced markedly high SARS-CoV-2-specific antibodies and induced a robust cellular immunity response compared to an electroporated DNA vaccine. mRNA-encoded antigens more closely resemble the structure and presentation of viral proteins expressed during a natural infection. mRNA vaccines have a vector-less approach and thus can avoid the potential for diminished immunogenicity with repeat dosing sometimes observed with vector-based vaccines. Moreover, the utility of intramuscularly administered mRNA vaccines against respiratory pathogens such as SARS-CoV-2 has been established, demonstrating robust immune responses and high real-world effectiveness against disease. However, the best effective LNPs for vaccination need to be uncovered and investigated further.

In the present study, we underline the crucial role of delivery systems, i.e., lipid nanoparticles in the efficacy of vaccines. LNPs improve the pharmacokinetics of the mRNA vaccine, such as its distribution and stability in the body, which can enhance the vaccine’s overall efficacy. LNPs need to be tailored with specific lipid compositions, which can maximize the effective immune response by promoting certain pathways of the immune system. Therefore, in this preclinical study, using SARS-CoV-2 as a model pathogen, the immunogenicity and protective efficacy of different LNPs encoding SARS-CoV-2 specific spike were evaluated. In this study, we expand upon the effect of LNP formulation on induction of immune responses. Studies have shown that there is an intrinsic ability of the LNPs to promote IL-6 secretion in mice and subsequent TFH induction [41]. Here, we show that one specific LNP formulation-LNP4 induced higher maturation and activation of DCs as measured by the frequency of co-stimulatory surface receptors and the production of cytokines and chemokines. By enhancing APC maturation and cytokine production, LNP-4 likely promotes a stronger and more durable immune response to the vaccine antigen compared to other LNPs. Additionally, a 2-dose primary i.m. vaccination regimen of mRNA (Spike)-LNP elicited systemic immune responses and resulted in lower SARS-CoV- 2 infection levels and disease severity versus mock-vaccinated controls after the viral challenge. Vaccination with mRNA-LNP4 showed improved protection against SARS-CoV-2. Based on the findings from this study, we have used LNP-4 while formulating all our mRNA-LNP based COVID-19 vaccines which we found to be highly efficient in providing antigenicity and immunity against multiple SARS-CoV-2 variants of concerns [42].

The biophysical parameters of LNP largely impacts its immunogenicity and an understanding of the same is an important step for enabling the rapid development of potent mRNA-LNP vaccines. Different mRNA vaccine formulations developed for protection against infectious conditions have entered clinical studies to evaluate their effectiveness. The size of LNPs for example can have an impact on their immunogenicity, as demonstrated by studies such as the one conducted by Hassett et al. and Brewer et al. [43] In the study by Hassett et al., the researchers investigated the effect of different biophysical factors on the immunogenicity of a cytomegalovirus (CMV) mRNA-LNP vaccine in mice. They found that LNP size had the strongest correlation with immunogenicity, with larger LNPs (up to 100 nm) leading to higher antibody titers. Similarly, Brewer et al. evaluated the antibody response to ovalbumin (OVA)-loaded liposomes of different sizes (100, 155, 225, and 560 nm) in mice [44]. They found that larger vesicles elicited a more significant IFN-γ response and higher levels of IgG2a, which is indicative of a Th1-type immune response. In contrast, smaller vesicles elicited higher levels of IgG1, which is indicative of a Th2-type immune response. In the present study, we observed that vaccination with mRNA encoding the spike protein of SARS-CoV-2 encapsulated in LNP-4 induces high levels of IgG antibodies and neutralizing antibodies against the virus. Additionally, peripheral blood obtained from Hamsters immunized with LNP-4 showed higher production of IFN-γ followed by hamsters immunized with other LNPs. LNP4 (IL00V41) comprised of DOPE (1,2-dioleoyl-sn-glycero-3-phosphoethanolamine), and the amount ionizable lipid used in the formulations was “higher” for LNP4 (IL00V41). It is hypothesized that the PEG-lipid is located only at the LNP surface, hence raising the mol% of the PEG-lipid (surface molecule) leads to a higher surface area: volume ratio and thus decrease in particle size. In our study, LNP4 displayed higher mol% range indicating that the effect of concurrently modifying cholesterol ratios to change the molar ratio of PEG, increases the immune efficacy of LNPs. Because of the same antigens (Spike) used, same dosing/immunization regimen, we establish the protective efficacy of various vaccine formulations. However, the mechanism of action of LNPs were not examined in these studies. Nevertheless, our study points out the importance of development and screening/testing of LNP formulations for favorable immunostimulatory profiles in studies such as these. The size and phospholipid dependence of LNPs for their efficacy are areas of ongoing research, with many aspects not fully understood. The role of specific phospholipid mixtures and off-target effects are yet not fully understood. Future studies will address these important questions.

## ACKNOWLEDGEMENTS

Studies of this report were supported by Public Health Service Research grants AI158060, AI150091, AI143348, AI147499, AI143326, AI138764, AI124911, and AI110902 from the National Institutes of Allergy and Infectious Diseases (NIAID) to LBM.

## REFERENCES

1. Janeway, C. and C. Janeway, Immunobiology : the immune system in health and disease. 5th ed. 2001, New York: Garland Pub. xviii, 732 pages : illustrations.

2. Quadiri, A., et al., Identification and characterization of protective CD8(+) T-epitopes in a malaria vaccine candidate SLTRiP. Immun Inflamm Dis, 2020. 8(1): p. 50–61.

3. Kashif, M., A. Quadiri, and A.P. Singh, Essential role of a Plasmodium berghei heat shock protein (PBANKA_0938300) in gametocyte development. Sci Rep, 2021. 11(1): p. 23640.

4. WHO, 2023 *WHO Global Vaccine Market Report*, in 2023 *WHO Global Vaccine Market Report*, WHO, Editor. 2023.

5. Quadiri, A., et al., SLTRiP induces long lasting and protective T-cell memory response. bioRxiv, 2021: p. 2021.01.07.425694.

6. Quadiri, A., et al., Therapeutic prime/pull vaccination of HSV-2-infected guinea pigs with the ribonucleotide reductase 2 (RR2) protein and CXCL11 chemokine boosts antiviral local tissue-resident and effector memory CD4(+) and CD8(+) T cells and protects against recurrent genital herpes. J Virol, 2024. 98(5): p. e0159623.

7. Belongia, E.A. and A.L. Naleway, Smallpox vaccine: the good, the bad, and the ugly. Clin Med Res, 2003. 1(2): p. 87–92.

8. Rodrigues, C.M.C. and S.A. Plotkin, Impact of Vaccines; Health, Economic and Social Perspectives. Front Microbiol, 2020. 11: p. 1526.

9. Mohammad, K., et al., Optimized plasmid loading of human erythrocytes for Plasmodium falciparum DNA transfections. Int J Parasitol, 2024.

10. Zayou, L., et al., A multi-epitope/CXCL11 prime/pull coronavirus mucosal vaccine boosts the frequency and the function of lung-resident memory CD4(+) and CD8(+) T cells and enhanced protection against COVID-19-like symptoms and death caused by SARS-CoV-2 infection. J Virol, 2023. 97(12): p. e0109623.

11. Rouzine, I.M. and G. Rozhnova, Evolutionary implications of SARS-CoV-2 vaccination for the future design of vaccination strategies. Commun Med (Lond), 2023. 3(1): p. 86.

12. Kalia, I., et al., Plasmodium berghei-Released Factor, PbTIP, Modulates the Host Innate Immune Responses. Front Immunol, 2021. 12: p. 699887.

13. Dhanushkodi, N.R., et al., Antiviral and Anti-Inflammatory Therapeutic Effect of RAGE- Ig Protein against Multiple SARS-CoV-2 Variants of Concern Demonstrated in K18-hACE2 Mouse and Syrian Golden Hamster Models. J Immunol, 2024. 212(4): p. 576–585.

14. DeFrancesco, L., Whither COVID-19 vaccines? Nat Biotechnol, 2020. 38(10): p. 1132–1145.

15. Hou, X., et al., Lipid nanoparticles for mRNA delivery. Nat Rev Mater, 2021. 6(12): p. 1078–1094.

16. Swetha, K., et al., Recent Advances in the Lipid Nanoparticle-Mediated Delivery of mRNA Vaccines. Vaccines (Basel), 2023. 11(3).

17. Reichmuth, A.M., et al., mRNA vaccine delivery using lipid nanoparticles. Ther Deliv, 2016. 7(5): p. 319–34.

18. Wilson, B. and K.M. Geetha, Lipid nanoparticles in the development of mRNA vaccines for COVID-19. J Drug Deliv Sci Technol, 2022. 74: p. 103553.

19. Tilstra, G., et al., Iterative Design of Ionizable Lipids for Intramuscular mRNA Delivery. J Am Chem Soc, 2023. 145(4): p. 2294–2304.

20. Lou, G., et al., Delivery of self-amplifying mRNA vaccines by cationic lipid nanoparticles: The impact of cationic lipid selection. J Control Release, 2020. 325: p. 370–379.

21. Kiaie, S.H., et al., Recent advances in mRNA-LNP therapeutics: immunological and pharmacological aspects. J Nanobiotechnology, 2022. 20(1): p. 276.

22. Li, Y., et al., A Comprehensive Review of the Global Efforts on COVID-19 Vaccine Development. ACS Cent Sci, 2021. 7(4): p. 512–533.

23. Zhou, L.Y., et al., Current RNA-based Therapeutics in Clinical Trials. Curr Gene Ther, 2019. 19(3): p. 172–196.

24. Saadati, F., S. Cammarone, and M.A. Ciufolini, A Route to Lipid ALC-0315: a Key Component of a COVID-19 mRNA Vaccine. Chemistry, 2022. 28(48): p. e202200906.

25. Jyotsana, N., et al., Lipid nanoparticle-mediated siRNA delivery for safe targeting of human CML in vivo. Ann Hematol, 2019. 98(8): p. 1905–1918.

26. Hald Albertsen, C., et al., The role of lipid components in lipid nanoparticles for vaccines and gene therapy. Adv Drug Deliv Rev, 2022. 188: p. 114416.

27. Anderluzzi, G., et al., Investigating the Impact of Delivery System Design on the Efficacy of Self-Amplifying RNA Vaccines. Vaccines (Basel), 2020. 8(2).

28. Nakhaei, P., et al., Liposomes: Structure, Biomedical Applications, and Stability Parameters With Emphasis on Cholesterol. Front Bioeng Biotechnol, 2021. 9: p. 705886.

29. Bailey, A.L. and P.R. Cullis, Membrane fusion with cationic liposomes: effects of target membrane lipid composition. Biochemistry, 1997. 36(7): p. 1628–34.

30. Heyes, J., et al., Cationic lipid saturation influences intracellular delivery of encapsulated nucleic acids. J Control Release, 2005. 107(2): p. 276–87.

31. Tenchov, R., et al., Lipid Nanoparticles horizontal line From Liposomes to mRNA Vaccine Delivery, a Landscape of Research Diversity and Advancement. ACS Nano, 2021. 15(11): p. 16982–17015.

32. Lin, P.J., et al., Influence of cationic lipid composition on uptake and intracellular processing of lipid nanoparticle formulations of siRNA. Nanomedicine, 2013. 9(2): p. 233–46.

33. Quadiri, A., et al., Antibody Responses Against Plasmodium falciparum MSP3 Protein During Natural Malaria Infection in Individuals Living in Malaria-Endemic Regions of India. Proc Natl Acad Sci India Sect B Biol Sci, 2022. 92(3): p. 613–619.

34. Prakash, S., et al., Cross-protection induced by highly conserved human B, CD4(+), and CD8(+) T-cell epitopes-based vaccine against severe infection, disease, and death caused by multiple SARS-CoV-2 variants of concern. Front Immunol, 2024. **15**: p. 1328905.

35. Xie, X., et al., Neutralization of SARS-CoV-2 spike 69/70 deletion, E484K and N501Y variants by BNT162b2 vaccine-elicited sera. Nat Med, 2021. 27(4): p. 620-621.

36. Muruato, A.E., et al., A high-throughput neutralizing antibody assay for COVID-19 diagnosis and vaccine evaluation. Nat Commun, 2020. 11(1): p. 4059.

37. Dhanushkodi, N.R., et al., Mucosal CCL28 Chemokine Improves Protection against Genital Herpes through Mobilization of Antiviral Effector Memory CCR10+CD44+ CD62L-CD8+ T Cells and Memory CCR10+B220+CD27+ B Cells into the Infected Vaginal Mucosa. J Immunol, 2023. 211(1):p. 118-129.

38. Prakash, S., et al., Genome-Wide B Cell, CD4(+), and CD8(+) T Cell Epitopes That Are Highly Conserved between Human and Animal Coronaviruses, Identified from SARS-CoV-2 as Targets for Preemptive Pan-Coronavirus Vaccines. J Immunol, 2021. 206(11): p. 2566-2582.

39. Wu, Z. and T. Li, Nanoparticle-Mediated Cytoplasmic Delivery of Messenger RNA Vaccines: Challenges and Future Perspectives. Pharm Res, 2021. 38(3): p. 473–478.

40. Chaudhuri, A., et al., Lipid-Based Nanoparticles as a Pivotal Delivery Approach in Triple Negative Breast Cancer (TNBC) Therapy. Int J Mol Sci, 2022. 23(17).

41. Alameh, M.G., et al., Lipid nanoparticles enhance the efficacy of mRNA and protein subunit vaccines by inducing robust T follicular helper cell and humoral responses. Immunity, 2021. 54(12): p. 2877–2892 e7.

42. Prakash, S., et al., A Broad-Spectrum Multi-Antigen mRNA/LNP-Based Pan- Coronavirus Vaccine Induced Potent Cross-Protective Immunity Against Infection and Disease Caused by Highly Pathogenic and Heavily Spike-Mutated SARS-CoV-2 Variants of Concern in the Syrian Hamster Model. bioRxiv, 2024.

43. Hassett, K.J., et al., Impact of lipid nanoparticle size on mRNA vaccine immunogenicity. J Control Release, 2021. 335: p. 237–246.

44. Watson, D.S., A.N. Endsley, and L. Huang, Design considerations for liposomal vaccines: influence of formulation parameters on antibody and cell-mediated immune responses to liposome associated antigens. Vaccine, 2012. 30(13): p. 2256–72.

