## Supplementary material for "Spike mRNA Vaccine Encapsulated in a Lipid Nanoparticle Composed of Phospholipid 1,2-Dioleoyl-sn-Glycero-3-Phosphoethanolamine Induced Potent B- and T-cell Responses Associated with Protection against COVID-19 in Hamsters": PDF

**Table 1: Physical Properties associated with the LNPs**

| LNP | Precision Nanosystems Nomenclature | Particle size (d.nm) | PDI | Encapsulation efficiency (%) | Mol % IL |
|---|---|---|---|---|---|
| LNP-1 | IL00V02 | 96 | 0.165 | 97.7 | Higher |
| LNP-2 | IL00V03 | 101 | 0.158 | 97.3 | Higher |
| LNP-3 | IL00V29 | 87 | 0.160 | 97.6 | Lower |
| LNP-4 | IL00V41 | 94 | 0.154 | 98.4 | Higher |

LNP: Lipid Nano Particle; PDI: Polydispersity index; Mol%: Relative amount of ionizable lipid present in the lipid mixture; IL: Ionizable lipid
